# The cellular esterase FrmB controls metabolic homeostasis and small colony variant formation in *Staphylococcus aureus*

**DOI:** 10.64898/2026.03.26.714663

**Authors:** Kelsey A O’Brien, Justin J Miller, R. Jeremy Johnson, Geoffrey C Hoops, Audrey R Odom John

**Author notes:** corresponding author Email: Audrey R. Odom John.

## Abstract

*Staphylococcus aureus* is a gram-positive bacterium that commonly colonizes the nasal passage, axilla, groin, and gastrointestinal tract of adults and adolescents. Although frequently nonpathogenic, *S. aureus* can infect most tissue types with clinical outcomes ranging from mild to fatal. The ability of *S. aureus * to persist and infect multiple body sites is driven largely by its metabolism. *S. aureus* can metabolize a wide range of niche-specific carbon sources, which contributes significantly to its persistence. Specifically, the utilization of glycolytic intermediates and pyruvate are associated with *S. aureus* bacterial burden and survival in the host. In this study, we establish the biological role of a serine hydrolase, FrmB, and evaluate its impact on *S. aureus* carbon metabolism. Using targeted metabolomics on *S. aureus* strains with and without FrmB, we find that FrmB is required for central carbon metabolic homeostasis. Mechanistically, we find that FrmB controls the enzymatic activity of the crucial rate-limiting enzyme, pyruvate dehydrogenase, which links glycolysis to the downstream tricarboxylic acid (TCA) cycle. Importantly, we find that the metabolic derangements in the absence of FrmB impact the ability of *S. aureus* to utilize pyruvate as a primary carbon source and reduce fitness. Finally, we demonstrate that FrmB is important for the formation of small colony variants (SCVs), a clinically relevant metabolic transition that is associated with chronic infection and antibiotic resistance.

## INTRODUCTION

*Staphylococcus aureus* is a gram-positive bacterium that is a common colonizer of the nasal passage, axilla, groin, and gastrointestinal tract of adults and adolescents^1–3,5^. Although frequently nonpathogenic, *S. aureus* often causes infection in both hospital and community settings. Between 15-50% of humans are persistently colonized by *S. aureus,* highlighting the size of the community reservoir and potential infection risk. ^1–3,5^ *S. aureus* can elicit infection across most tissue types with disease ranging from mild to fatal, and clinical manifestations include skin and soft tissue infections, staphylococcal endocarditis, staphylococcal osteomyelitis, and bacteremia. In the United States, methicillin-resistant *S. aureus* (MRSA) causes over 300,000 infections and over 10,000 deaths annually, highlighting the need for novel therapeutics and an enhanced understanding of staphylococcal virulence factors.^4^

In several bacterial species, the transition from colonization to invasive disease requires a rewiring of metabolic processes to facilitate effective use of site-specific nutrients.^1,2^ For example, the metabolism of *Streptococcus mutants* has evolved to persist in the modern human oral cavity with the shift towards carbohydrate heavy diets. This organism has evolved to contain additional carbohydrate metabolism genes that increase its capacity for carbohydrate-catabolite repression, a process which allows bacteria to select which carbohydrates to use for metabolism by sensing available extracellular nutrient content.^6–8^ Because *S. aureus* can infect most human organs and tissue types, its metabolism is expected to be both dynamic and integral to virulence, especially with respect to carbon metabolism.

*S. aureus* can utilize more than 15 carbohydrates as its primary carbon source. Of these, *S. aureus* favors glucose, which is converted by staphylococci into pyruvate and subsequently acetyl coenzyme A (acetyl-CoA).^9–10^ Unlike other organisms, *S. aureus* does not funnel all acetyl-CoA directly into the tricarboxylic acid (TCA) cycle (also known as the Krebs cycle or citric acid cycle) during aerobic, exponential phase growth. Even when extracellular glucose is abundant, acetyl-CoA is preferentially converted to acetate for extracellular export, a process known as overflow metabolism, allowing the cell to generate two additional ATP molecules per glucose.^11^ When environmental glucose is depleted, the exported acetate is imported back into the cell, converted into acetyl-CoA, and then funneled into the TCA cycle.^11–13^ This process is mediated by carbon (glucose) catabolite repression of proteins responsible for regulation of TCA cycle-associated enzymes.^14^

Central carbon metabolism of *S. aureus* has been clearly linked to disease states and niche occupations. For example, intracellular glucose-6-phosphate levels are associated with induction of staphylococcal cytotoxins, which mediate tissue necrosis and overall bacterial burden.^15^ Similarly, extracellular glucose and acetate impact *S. aureus* virulence in a diabetic mouse model, although other studies demonstrate that excessive levels of extracellular glucose represses toxin production and reduce virulence.^11,16–18^

Perhaps the most studied clinical occurrence of a central carbon metabolism-dependent adaptation is the transition of *S. aureus* to the small colony variant form. On solid media, small colony variants (SCVs) appear as slow growing, petite colonies, and, in the clinical setting, SCVs are associated with chronic infection, antibiotic resistance, and host persistence.^19–20^ When transitioning to a SCV form, *S. aureus* rewires its metabolism to repress the TCA cycle and electron-transport chain.^21^ To compensate, SCV staphylococci upregulate glycolysis, as well as pyruvate, butanoate, arginine, and nitrate metabolism.^21–22^ While more metabolically quiescent, SCVs decrease the expression of major virulence regulators, allowing the bacteria to evade host immune system detection. Additionally, the decrease in SCV oxygen consumption results in the bacteria failing to activate host-oxygen sensing proteins (HIF-1) that alert intracellular invasion has occurred, contributing to persistence in the host.^23^ This decrease in cellular respiration also confers resistance to antibiotics, such as aminoglycosides.^24^ As a result, SCVs are associated with worse clinical outcomes and represent a significant challenge with treating chronically infected patients, further highlighting the need understand metabolic regulation in this pathogen.^19^

Our lab has recently identified and enzymatically and structurally characterized a novel serine-hydrolase in *S. aureus*, FrmB.^25^ This enzyme demonstrates high structural conservation to esterases in *S. pneumonia* and *B. intestinalis* that are involved in virulence and ferulic acid metabolism respectively.^25–27^ Additionally, we find that FrmB is highly conserved across *S. aureus* phylogeny, suggesting an important function in growth or persistence.^25^ Here, we perform targeted metabolomics on *S. aureus* with and without FrmB and discover that FrmB is important for homeostasis of central carbon metabolism. We further demonstrate that FrmB is required for optimal function of pyruvate dehydrogenase and that staphylococci require FrmB to effectively utilize pyruvate as a carbon source. Finally, metabolic defects in cells lacking FrmB lead to a defect in small colony variant formation. Our research establishes FrmB as a novel regulator of central carbon metabolism in *Staphylococcus aureus*, offering insights into the dynamic metabolism of this pathogen.

### FrmB is required for central carbon metabolic homeostasis in *S. aureus*

In our previous studies to evaluate prodrug activation in staphylococci, we identified and enzymatically characterized a serine hydrolase of unknown biological function, FrmB (SAUSA300_2564). This hydrolase, which can de-esterify intracellular prodrugs, has high structural homology to a suite of metabolic esterases from divergent species. Additionally, FrmB is present in all publicly available *S. aureus* sequences and demonstrates moderate conservation across all clonal complexes and high conservation within individual clonal complexes.^25^ We hypothesized that FrmB may be involved in a conserved metabolic pathway in *S. aureus*, due to its high structural similarity to other metabolic esterases and within-species sequence conservation.

To evaluate the role of FrmB in *S. aureus* metabolism, we compared the metabolic profile of wild-type *S. aureus* (JE-2) to an isogenic strain with a transposon insertion in FrmB (JE-2 *frmB*::tn). Both strains were grown to mid-exponential phase and targeted metabolomics was performed using a large panel representing carbon metabolites (supplemental methods 1). In total, 309 biochemicals were identified, 259 of known identity and 50 of unknown structural identity. Following normalization to DNA content and log transformation, we performed a principal components analysis (PCA) to visualize the discriminatory impact of FrmB on the *S. aureus* metabolome. We find significant differences in overall metabolite profiles based on the presence or absence of FrmB (Figure1A). The impact of FrmB on *S. aureus* metabolite composition is further appreciated in a hierarchical clustering analysis. As with our PCA analysis, we see differential clustering of metabolite profiles (Z-score) between *S. aureus* strains with and without FrmB, confirming that FrmB is required for global metabolic homeostasis in *S. aureus* (Figure 1B).

**Figure 1.**
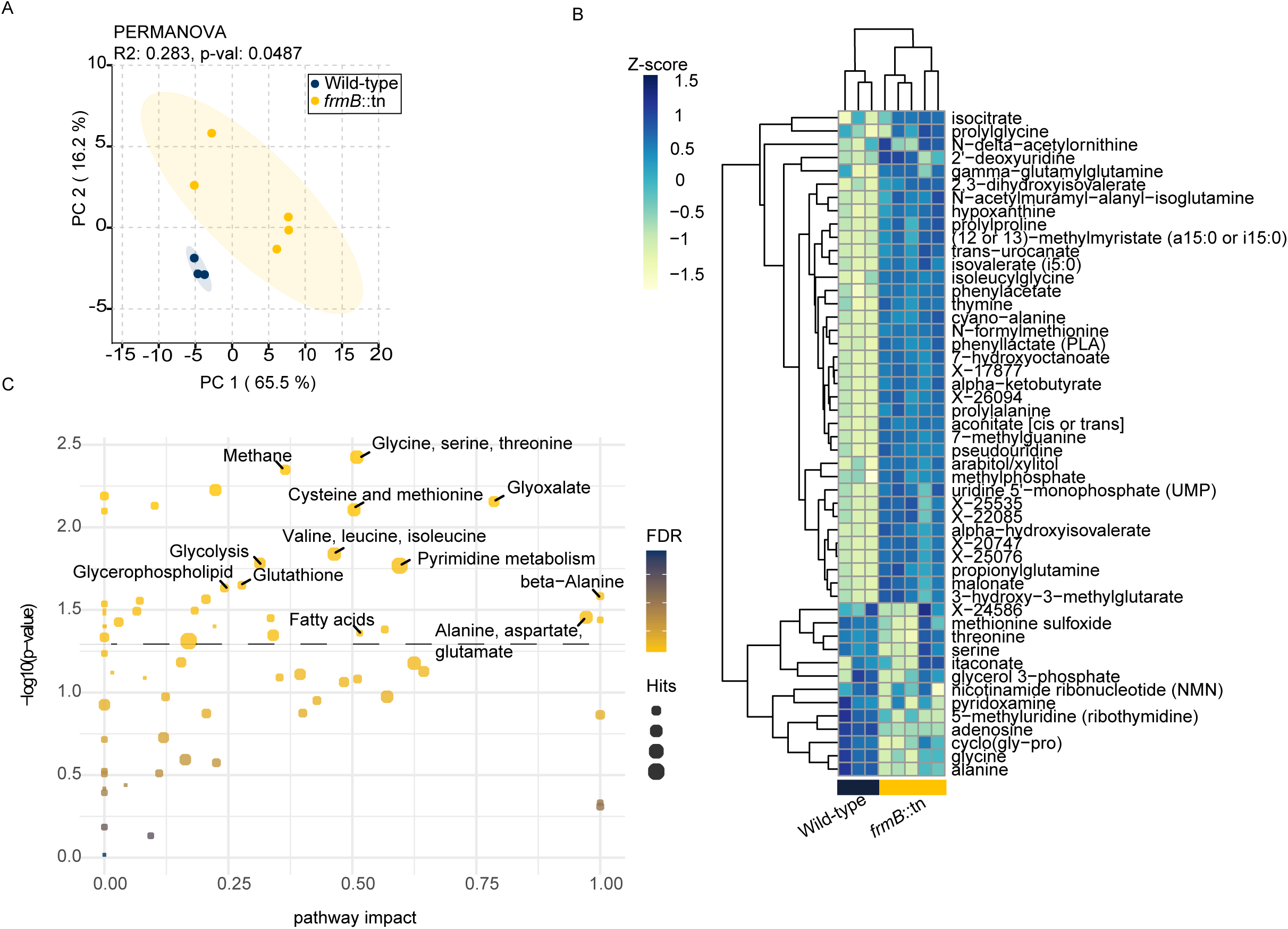
FrmB is required for normal metabolic homeostasis in *S. aureus*. A) PCA plot showing the discriminatory impact of FrmB on *S. aureus* metabolites; wild-type (blue), *frmB:*:tn (yellow) (PERMANOVA, R2: 0.283, P-val:0.0487). B) Hierarchical clustering analysis of the top 50 most impacted metabolites (Z-score) based on FrmB presence or absence (Ward clustering and Euclidean distance measured.) C) Pathway enrichment analysis performed using *Staphylococcus aureus* subs. Aureus USA300FPR3757 (CA-MRSA)(KEGG) metabolome as the reference. The pathway impact is reported against statistical significance. FDR is visualized by color, size of circle indicates number of metabolite hits that fit within the marked category.

To further characterize FrmB-dependent metabolic pathways, we performed a pathway enrichment analysis. We performed an ANCOVA analysis assessing the relative-betweenness centrality of metabolic pathways, using the *Staphylococcus aureus* subsp. *aureus* USA300FPR3757 (CA-MRSA) (KEGG) metabolome as a reference. We find that the most enriched metabolites are associated with pathways belonging to amino acid metabolism, glycolysis, and fatty acid synthesis (Figure 1C). Specifically, loss of FrmB leads to elevated steady-state levels of 22 metabolites and decreased levels of 20 metabolites. Serine, threonine, and leucine comprise the three most enriched metabolites, followed by additional amino acids, glucose, glucose 6-phosphate (G6P), dihydroxyacetone phosphate (DHAP), and pyruvate (Figure 2A, Supplemental figure 1A-C). Deoxyuridine, phosphoribosyl pyrophosphate (PRPP), two unidentified compounds, and, notably, acetyl-CoA were significantly decreased (Figure 2A, Supplemental figure 1C). Energetic markers adenosine triphosphate (ATP), adenosine diphosphate (ADP), and nicotinamine adenine dinucleotide (NAD+) were also decreased upon loss of FrmB (Supplemental figure 1E). We find that loss of FrmB is associated with a pattern of metabolic changes described by an increase in proximal glycolytic metabolites (and amino acids upstream of pyruvate), and a decrease in acetyl-CoA. In addition, loss of FrmB is also associated with decreased cellular levels of citrate, aconitate, isocitrate, succinate, malate, fumarate (Figure 2B, Supplemental figure 1D). Altogether, these data suggest that FrmB is required for metabolic homeostasis, and appears to set the balance of pyruvate versus acetyl-CoA, two metabolites that are typically interconverted by the enzyme pyruvate dehydrogenase.

**Figure 2.**
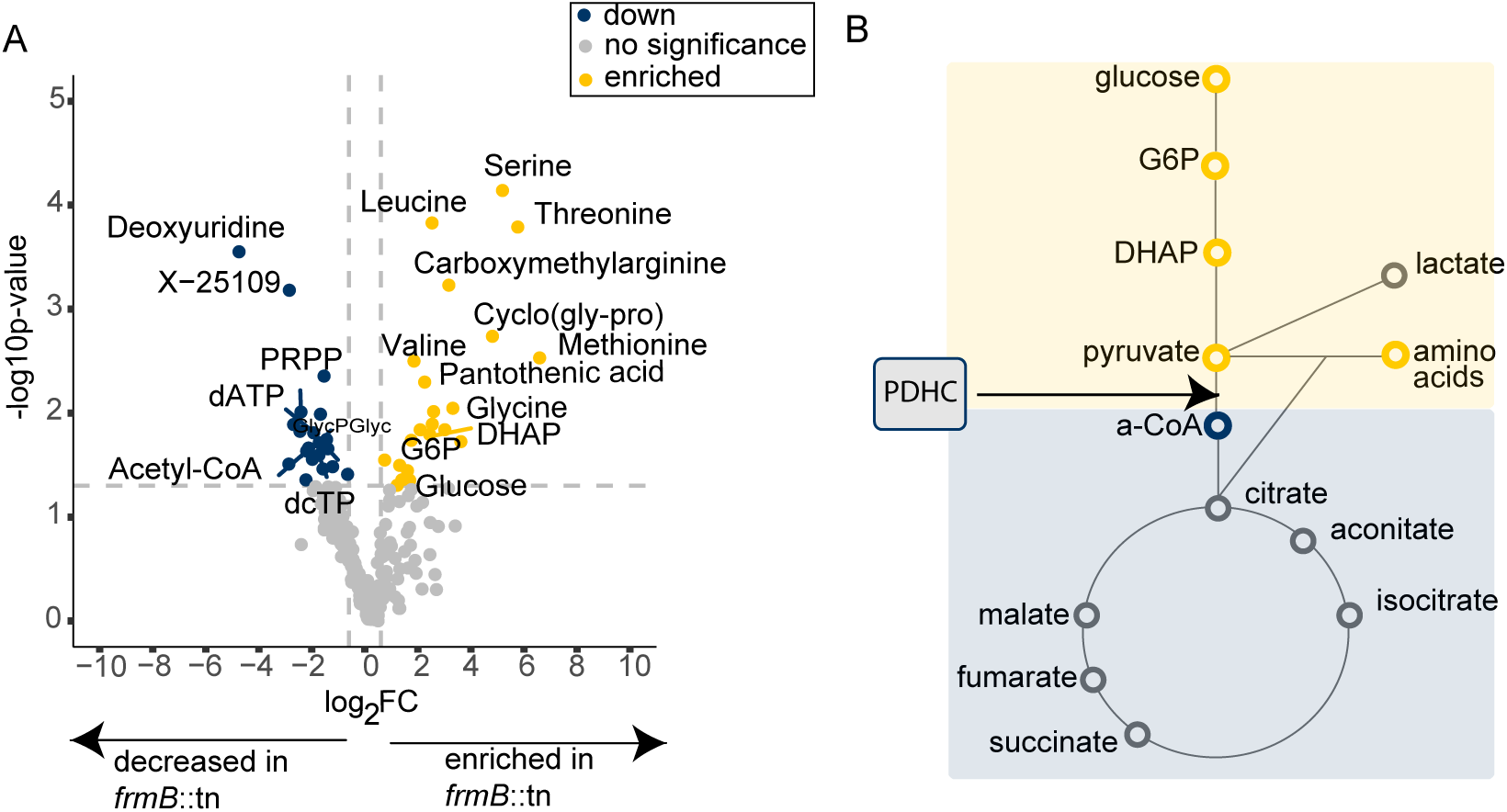
FrmB is required to maintain wild-type levels of proximal glycolytic metabolites and acetyl-coA. A) fold-change plot showing metabolites that are significantly enriched in *frmB:*:tn (yellow) and significantly decreased in *frmB:*:tn (blue.) A fold change cut-off of 2.0 was selected and a p-value threshold of 0.05 was applied. B) Schematic of select central carbon metabolites that were significantly enriched in *frmB:*:tn (yellow) or significantly decreased in in *frmB:*:tn (blue) or no significant change (gray).

### FrmB is required for basal cellular activity of pyruvate dehydrogenase

To confirm that phenotypes are due solely to loss of FrmB in the *frmB*::tn strain, we established that restoring FrmB expression would restore cellular function. Using a single-copy complementation strategy, we generated an FrmB complementation strain in the *frmB*::tn background (Figure 3A). We confirmed the genetic insertion via polymerase chain reaction (PCR) (Figure 3B) and confirmed that complementation restored cellular FrmB-dependent esterase activity, using our published fluorescent ester activation assay^25,28^. Out of 32 ester substrates, ester 6C is activated selectively by FrmB compared to other known *S. aureus* esterases, allowing us to establish FrmB specific activity restoration. We applied bacterial whole cell lysate from either wild-type, *frmB*::tn, or complement to ester 6C. De-esterification of this substrate causes increased fluorescence, proportional to enzymatic activity of FrmB (Figure 3C). As expected, the *frmB*::tn strain has markedly reduced FrmB esterase activity compared to wild-type, which is substantially improved by FrmB complementation, although it does not fully reach wild-type levels.

**Figure 3.**
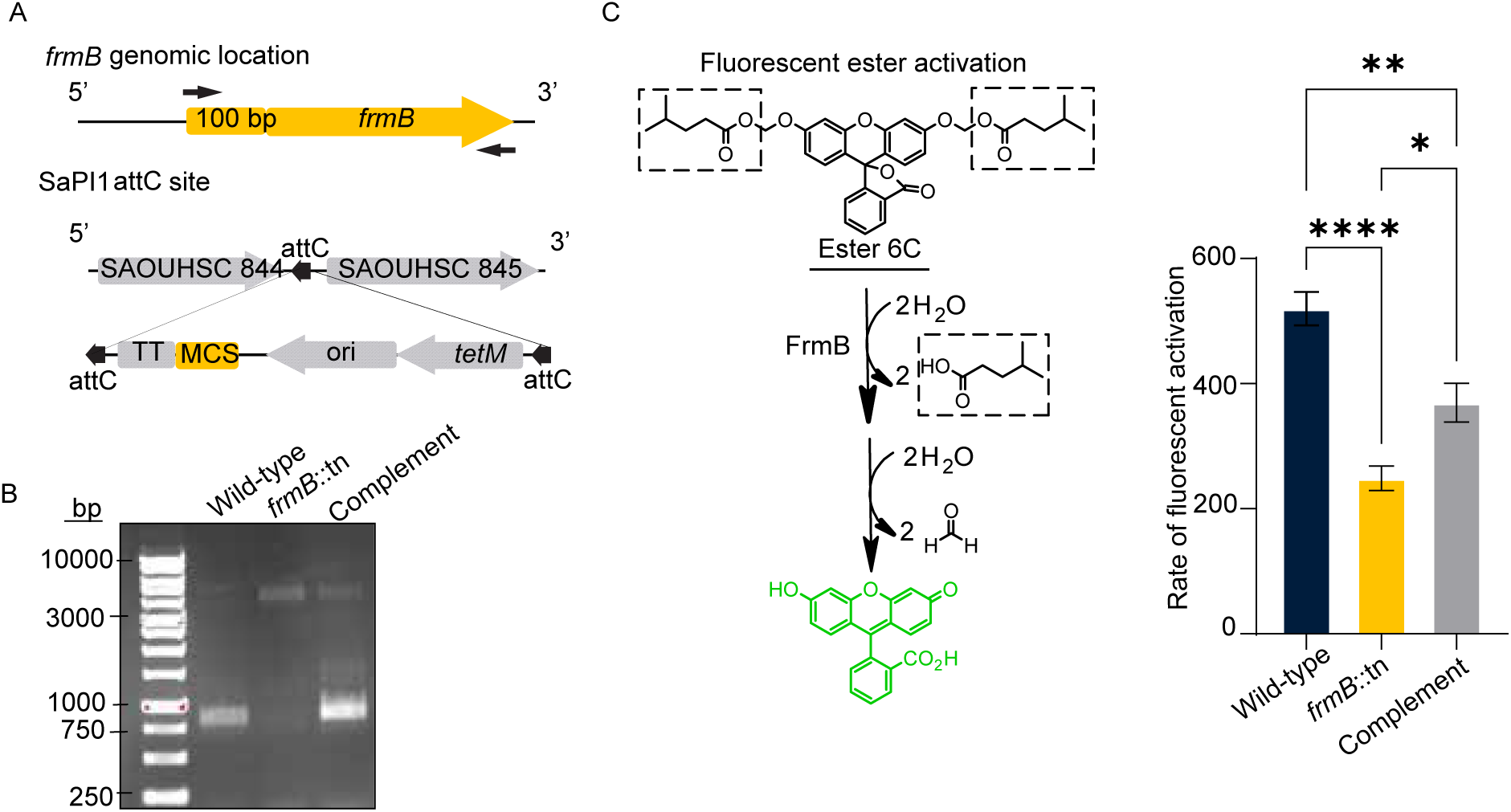
Complementing FrmB restores known FrmB enzymatic function . A) SaPI1 *S. aureus* single copy complementation protocol schematic. FrmB and 100bp upstream were cloned into the SaPI1 attC site in *frmB::*tn. B) PCR confirmation of *frmB* and upstream region integration into *frmB:*:tn. C) Fluorescent ester activation confirming native FrmB enzymatic activity. A known FrmB specific substrate (ester 6C) was added to wild-type, *frmB::*tn, or complement lysate and fluorescence was measured as a proxy for enzyme activity,error bars represent SEM (One-way ANOVA, Tukey’s correction for multiple comparisons, * P≤0.05, *** P ≤ 0.001, *** P≤ 0.0001.)

Because *S. aureus* cells lacking FrmB have an abnormal balance of pyruvate versus acetyl-CoA, we next sought to evaluate whether the presence or absence of FrmB impacts the cellular enzymatic activity of pyruvate dehydrogenase in *S. aureus*. The pyruvate dehydrogenase complex (PDHC) comprises three distinct subunits, each of which catalyzes a different step in the conversion of pyruvate into acetyl-CoA.^29^ We hypothesized that the FrmB-associated accumulation of upstream metabolites and decrease in acetyl-CoA was due to a reduction in the enzymatic activity of the E1-subunit of PDHC (PdhA). We therefore leveraged a PDHC activity assay that measures the rate at which the E1 subunit of PDHC can reduce an artificial electron acceptor, 2,6-dichlorophenolindophenol (DCPIP), when pyruvate is supplied as a substrate (Figure 4A)^30^. We cultured *S. aureus* with or without FrmB in complete media to mid-exponential phase, prepared lysates, and measured native PDHC activity. When FrmB is absent, there is a significant decrease in the rate of DCPIP reduction, but this reduction is restored by FrmB complementation. This suggests that FrmB is required for the basal activity of PDHC in *S. aureus* (Figure 4B).

**Figure 4.**
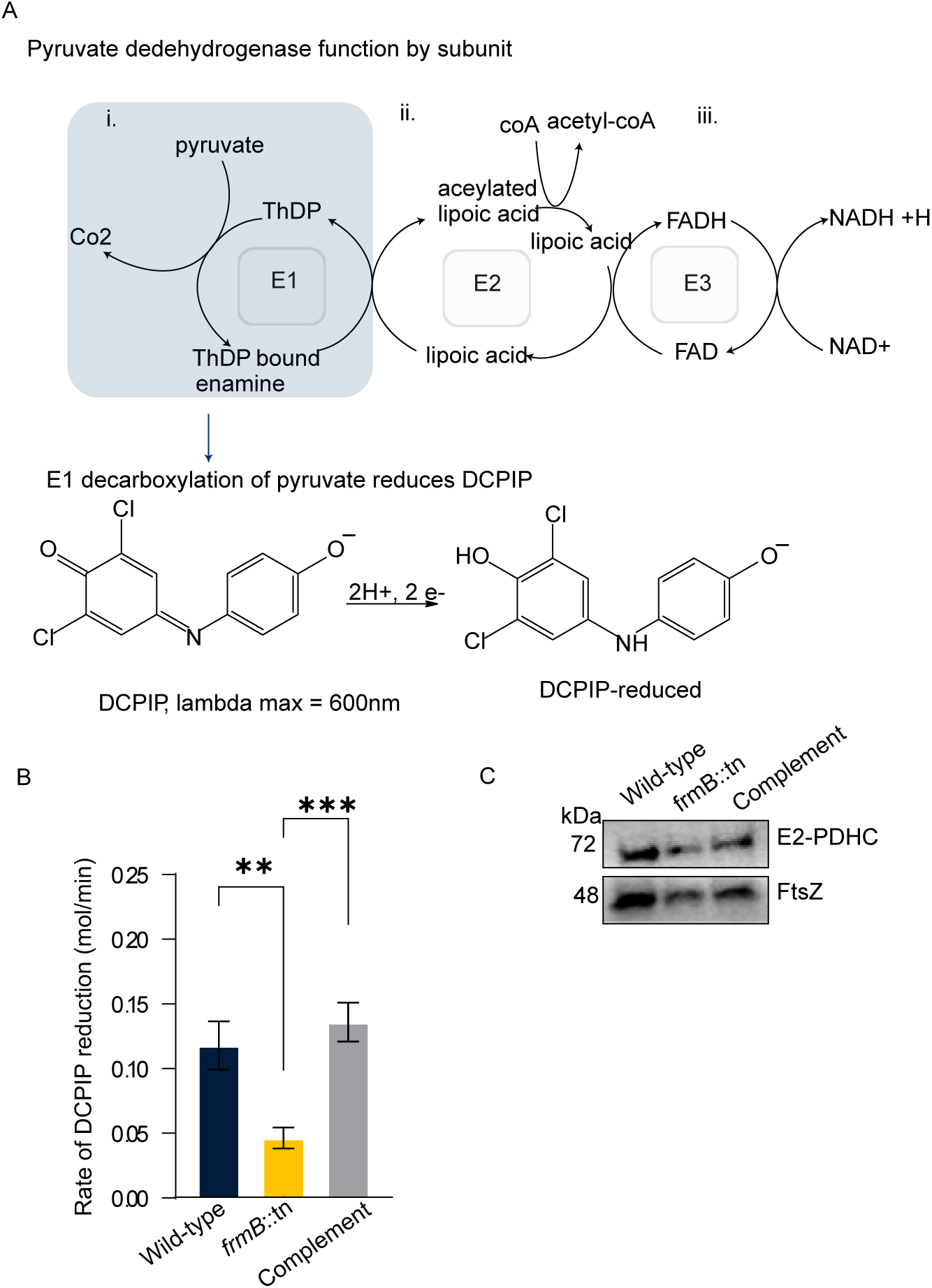
Loss of FrmB results in decreased E1-PDHC activity. A) 2,6-Dichlorophenolindophenol (DCPIP) was utilized as an electron donor to assess the activity of PDHC. Briefly, the E1 subunit of PDHC decarboxylates pyruvate, which facilitates a two electron reduction of DCPIP that can be monitored overtime by a change in absorbance. B) rate of DCPIP reduction for wild-type, *frmB::*tn, and complement whole cell lysates. Shown here are three technical replicates representative of three independent biological replicates, error bars represent SEM. (One-Way ANOVA, Tukey’s correction for multiple comparisons, * P≤0.05, *** P ≤ 0.001, *** P≤ 0.0001.) C) Western blot showing PDHC levels by probing the E2 subunit of PDHC via anti-lipoic acid antibody. An anti-FtsZ (essential S. aureus cell division protein) antibody was utilized as a loading control. Shown here is a representative blot from three biological replicates.

To establish if this phenotype was due to reduced abundance of PDHC protein when FrmB is absent, we quantified PDHC protein levels via immunoblot. We observed no differences in PDHC protein levels, suggesting that the observed reduction in PDHC activity in cells lacking FrmB is not due to reduced expression (Figure 4C). Together, these data suggest that FrmB impacts the catalytic activity, but not expression, of PDHC.

### FrmB is required for efficient growth under carbon-limited conditions

Because our data suggest that FrmB impacts central carbon metabolism at the level of pyruvate dehydrogenase, we predicted that FrmB might likewise impact growth and/or fitness of *S. aureus*. In carbon-replete rich media, we found no difference in growth based on the presence or absence of FrmB (Figure 5A). However, in competition growth assays in which strains with and without FrmB were grown together, we observed that wild-type *S. aureus* out-competed the mutant lacking FrmB over the course of 24 hours, suggesting that FrmB plays a significant role in *S. aureus* fitness (Figure 5C).

**Figure 5.**
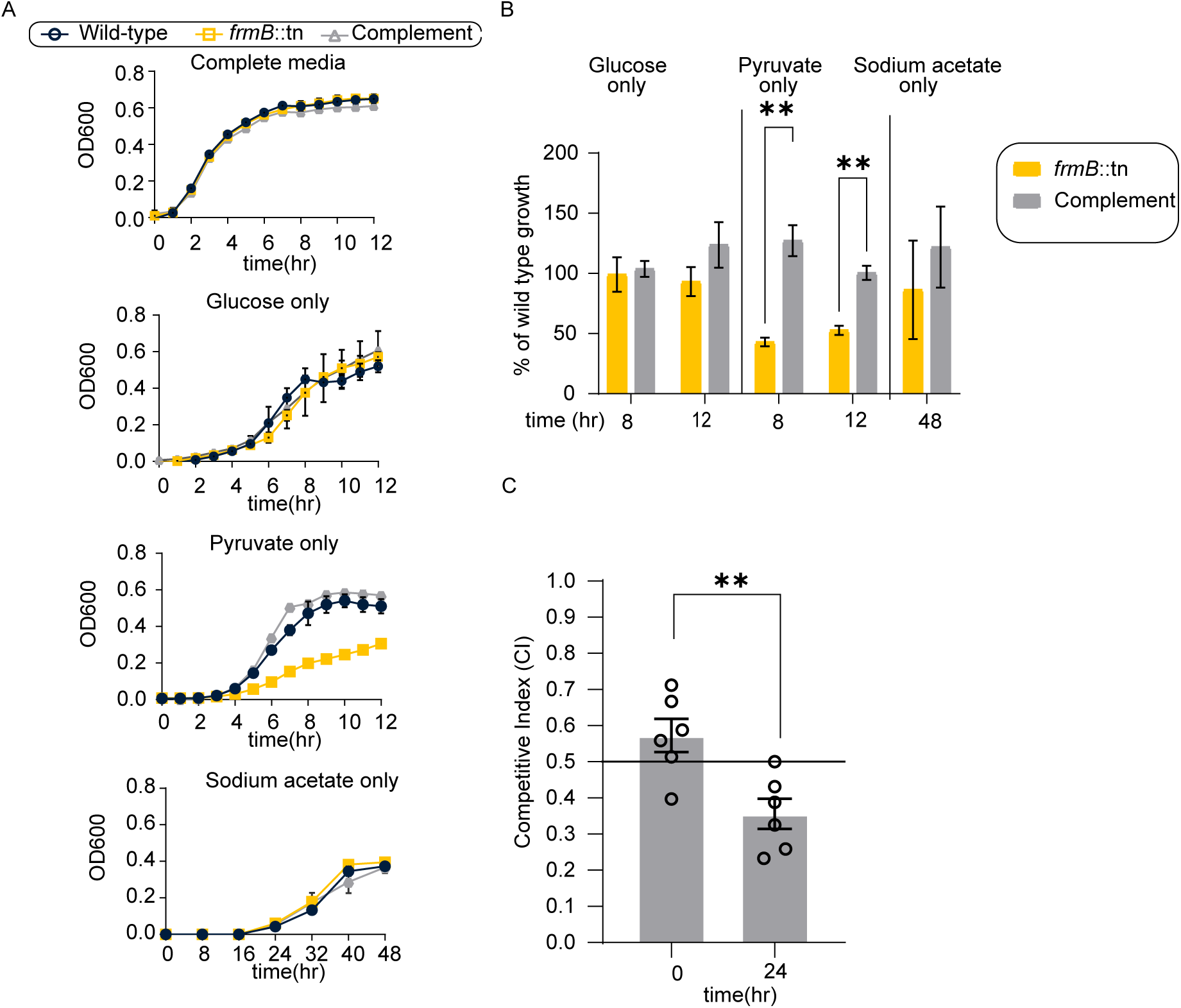
Loss of FrmB results in growth attenuation under carbon scarce conditions. A) Representative growth of wild-type, *frmB:*:tn, and complement in complete media and drop out media. The sole carbon source added to support growth is listed above each curve, error bars represent SEM of technical replicates. B) means and SEM for three experimental replicates at 8 hours, 12 hours, or 24 hours of growth for each carbon source are shown. (multiple unpaired t-tests, FDR, two stage step-up Benjamini, Krieger, and Yekutieli, * P≤0.05, *** P ≤ 0.001, *** P≤ 0.0001.) C) competitive index of wild-type and frmB::tn in rich media, means and SEMs from independent experiments are reported (Welch’s t-test, * P≤0.05, *** P ≤ 0.001, *** P≤ 0.0001.) Competitive index was calculated by dividing total CFUs on *frmB::*tn selective plates (LB/erm) by the total CFUs on plain LB. A competitive index < 0.5 indicates wild-type out competes *frmB:*:tn.

While loss of FrmB does not confer a growth defect in a rich media monoculture, our rich media contains glucose as the primary carbon source. Because FrmB impacts PDHC activity, we queried whether reduced conversion of pyruvate to acetyl-CoA would correlate with impaired growth when pyruvate was provided as a sole carbon source. We therefore evaluated growth in a defined minimal media that lacked a major carbon source, but contained trace amino acids necessary to support *S. aureus* growth (Supplemental table 2). As expected, *S. aureus* does not grow in the absence of exogenous carbon (Supplemental figure 2). Using this media, we added back either glucose, pyruvate, or sodium acetate and assessed the ability of our isolates to grow. In the absence of FrmB, we observed no growth defect when glucose was supplied as the major carbon source; however, when pyruvate was the sole carbon source provided, there was a significant attenuation in growth, which was restored upon genetic complementation (figure 5A,B). During exponential phase and aerobic glycolysis, *S. aureus* preferentially converts acetyl-CoA into acetate and subsequently exports this acetate from the cell rather than funneling acetyl-CoA into the TCA cycle (Supplemental figure 3).^13^ *S. aureus* can also use acetate as a sole carbon source when no other major sources are present, importing free acetate back into the cell. This imported acetate is converted directly into acetyl-CoA, thus bypassing the need for PDHC (supplemental figure 3).^13^ We therefore predicted that FrmB would not be required for acetate-dependent growth. Indeed, we observed no difference in growth when sodium acetate was provided as the sole carbon source (figure 5A,B).

Because FrmB is required for pyruvate dehydrogenase activity and growth with pyruvate as a carbon source, we hypothesized that FrmB would likewise impact overall glycolytic efficiency. The Agilent seahorse instrumentation quantifies glycolytic flux based on extracellular acidification, the result of exported products of bacterial fermentation or glycolysis (such as lactate and acetate, respectively). Each strain was grown in rich media and washed and resuspended in carbon-free media. For each isolate, we measured extracellular acidification (ECAR) and oxygen consumption rate (OCR) as proxies for glycolysis and respiration respectively (Figure 6A). At t =12 minutes, glucose or pyruvate was injected into the wells, and the changes in ECAR and OCR were recorded. When pyruvate was added, there was a significant initial increase in media acidification when FrmB is present, and this increase was sustained for the duration of the assay, demonstrating that FrmB impacts glycolytic efficiency when pyruvate is supplied as a carbon source (Figure 6B-D). We saw a less pronounced, but similar trend for oxygen consumption. However, when glucose was added, we saw a significant increase in media acidification and oxygen consumption in wild-type *S. aureus*, but this phenotype was not rescued by complementation. This suggests that FrmB does not impact glycolytic flux when glucose is provided, perhaps due to alternate metabolic fates glucose may encounter upstream of PDHC. (Figure 6B-D). Taken together, our growth data and extracellular flux results suggest that FrmB contributes to the utilization of pyruvate in *S. aureus*.

**Figure 6.**
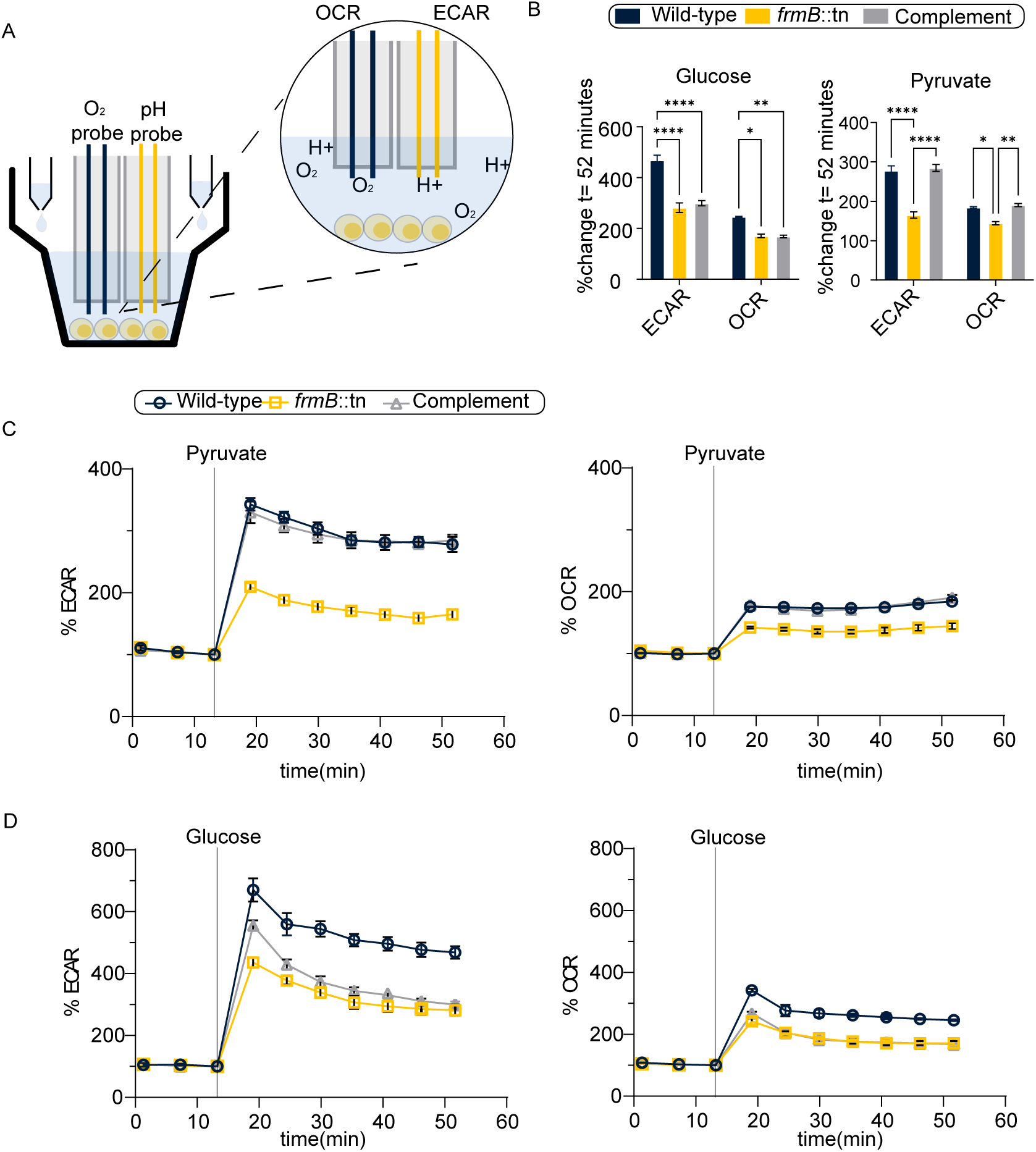
Loss of FrmB results in decreased glycolytic flux under carbon scarce conditions. A) Overview of seahorse extracellular flux set-up. In a 96-well plate, bacteria are seeded and changes in oxygen and pH are monitored overtime to capture extracellular acidification rate (ECAR) and oxygen consumption rate (OCR). Carbon sources are injected at specific timepoints via injection ports. B) Final changes in percent ECAR or percent OCR values for all biological replicates at the time of assay completion (t = 52 minutes), means and SEMs are reported (two-way ANOVA, Tukey’s correction for multiple comparisons, * P≤0.05, *** P ≤ 0.001, *** P≤ 0.0001, **** P≤ 0.00001.) Representative curves for changes in percent ECAR and percent OCR at the time of assay completion (t= 52 minutes) upon C) pyruvate addition or D) glucose addition for JE-2, *frmB::*tn, and complement. Carbon source injection is denoted by grey line. Means and error bars representing SEM of technical replicates are presented.

### FrmB is required for small colony variant formation

Treatment of staphylococcal infections can be challenging, as antibiotic therapy often fails to completely sterilize sites of infection, leading to recurrence once antibiotic pressure is removed. Infection recurrence is associated with the ability of *S. aureus* to form small colony variants (SCVs).^19–20^ Small colony variants are phenotypically distinct pin-point colonies with slow growth rates, which are clinically associated with enhanced antibiotic resistance and immune system evasion.^20^ Previous work has demonstrated that SCVs are marked by a unique metabolic rewiring that down-regulates the TCA cycle and respiration, and upregulates glycolysis, pyruvate metabolism, and alternative flux routes.^21–24^ Because FrmB impacts metabolic homeostasis at the level of pyruvate and lower glycolysis, we hypothesized that FrmB would also be required for this metabolic re-wiring and SCV production.

To test this hypothesis, we induced small colony variants through exposure to low pH over time.^30^ A low pH environment mimics the lysosome, which previous studies have demonstrated to be a reservoir of *S. aureus* SCVs in clinical settings.^31–32^ We cultured staphylococcal strains with and without FrmB in a defined, carbon-replete media. We acidified the media to pH 4.0 and allowed our isolates to persist statically at 37°C for 24 hours (Figure 7A). At 24 hours, we quantified viable bacteria (colony forming units) and formation of SCVs. We defined SCVs as pin-point colonies, which are visibly distinct and approximately one tenth the size of a normal colony (Figure 7C.)^31^ Consistent with previous literature, 24 hours of persistence at a pH of 4.0 successfully generated SCVs in wild-type staphylococci. We find that FrmB was not required for 24 hour survival at low pH (Supplemental figure 4). However, cells lacking FrmB were markedly impaired in SCV generation (Figure 7B).

**Figure 7.**
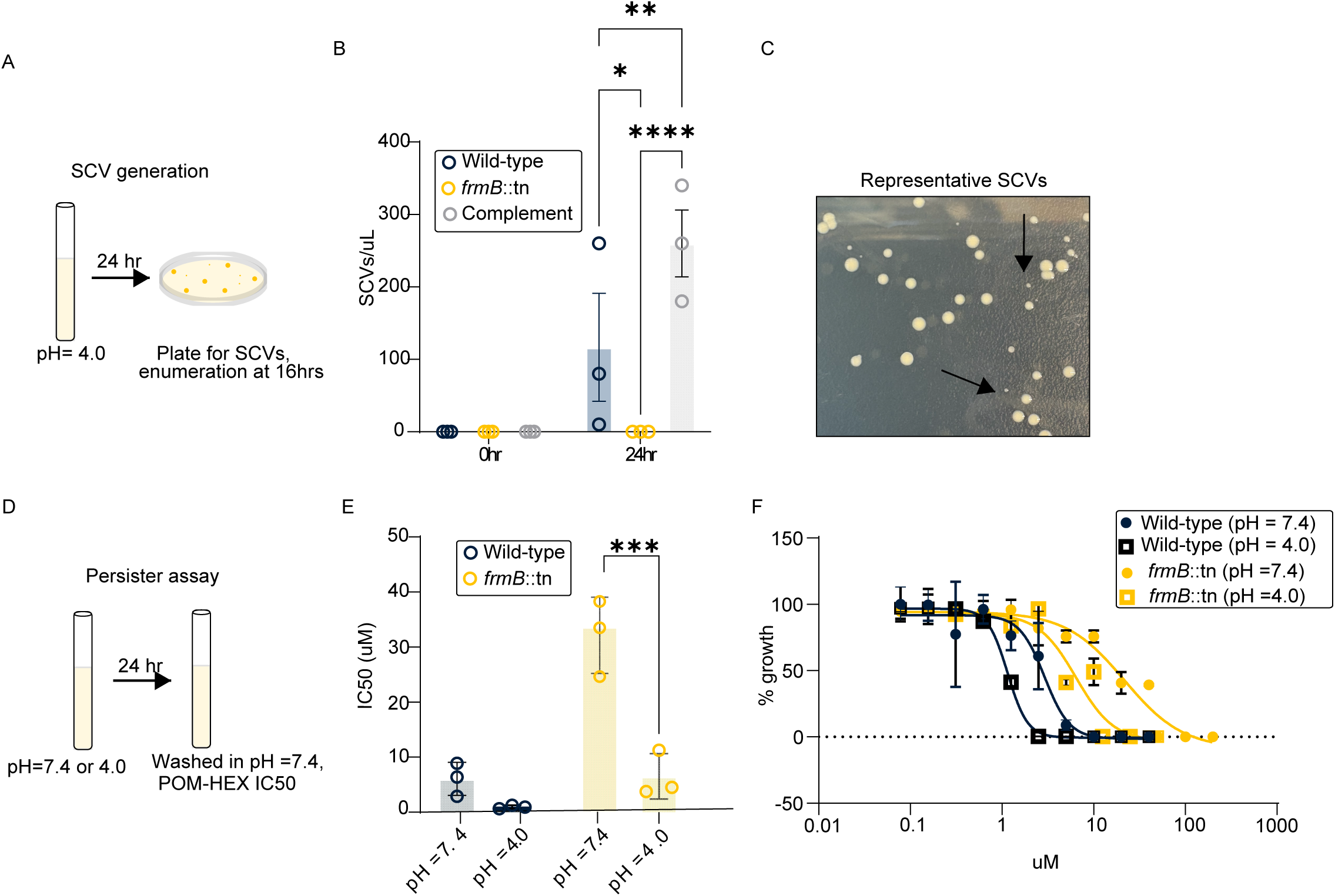
FrmB impacts small colony variant formation. A) Small colony variant (SCV) generation protocol. Isolates were exposed to a pH of 4.0 for 24-hours and B) SCVs were enumerated on LB plates as C) pin-point colonies (denoted by black arrows). Means and SEM are reported (two way ANOVA, Tukey’s correction for multiple comparisons, * P≤0.05, *** P ≤ 0.001, *** P≤ 0.0001.). D) Persister assay protocol. Isolates were exposed to a pH of 7.4 or a pH of 4.0 for 24 hours, washed in a pH of 7.4, then immediately exposed to POM-HEX. E) IC50 values from three independent experiments are reported with means and SEMs accompanied by F) representative IC50 curves (Two way-ANOVA Tukey’s correction for multiple comparisons, * P≤0.05, *** P ≤ 0.001, *** P≤ 0.0001.)

Finally, we have previously established that *S. aureus* FrmB is one of two esterases that can activate a prodrug, POM-HEX, which inhibits the glycolytic enzyme enolase.^25^ Resistance to POM-HEX can be acquired via spontaneous mutations in FrmB, resulting in a decreased ability of the bacteria to hydrolyze the carboxy ester promoities of the prodrug. Because FrmB impacts the ability to transition into a glycolytically dependent SCV state, we hypothesized that, when forced into said glycolytically dependent state, we would regain sensitivity to POM-HEX if FrmB is not present. Such that, the amount of prodrug activated by the second esterase will be sufficient to impair a glycolytically weakened mutant.

We utilized an established non-stable SCV persister assay.^33^ We exposed our isolates to a pH of 7.4 or 4.0 for 24 hours in a defined media. After 24 hours, we washed and resuspended each culture in a pH of 7.4 and immediately performed dose response assays for POM-HEX (Figure 7D). As expected, when grown at a pH pf 7.4, lacking FrmB confers a higher half-maximal inhibitory concentration (Figure 7E). However, after growth under SCV inducing conditions, both isolates are sensitized to POM-HEX, with *frmB*::tn having a half maximal inhibitory concentration comparable to wild-type under low stress conditions (Figure 7E). Our data demonstrate that, while a loss of FrmB confers resistance to POM-HEX under stress-free conditions, the fitness cost of losing FrmB under SCV-inducing conditions resensitizes the bacteria to this pro-drug.

In conclusion, our study demonstrates that *S. aureus* FrmB is a serine hydrolase involved in *S. aureus* central carbon metabolism. This esterase is responsible for the normal enzymatic activity of pyruvate dehydrogenase and this corresponds to the proper utilization of pyruvate as a carbon source and in metabolic flux. Additionally, when FrmB is not present, the defect accrued in pyruvate metabolism corresponds to a decreased ability to transition into a SCV state under low pH stress and a re-sensitization to an experimental antibacterial prodrug, demonstrating that FrmB is important for *S. aureus* persistence.

## DISCUSSION

The dynamic metabolism of *S. aureus* allows this pathogen to infect a myriad of host tissue types and is associated with chronic disease states and persistence in the host. Thus, understanding how *S. aureus* regulates and adapts its metabolism has profound implications to pathogenesis and potential new therapies. In this study, we sought to classify the biological role of a serine hydrolase, FrmB, and evaluate its impact on *S. aureus* metabolism. We find that FrmB is required for metabolic homeostasis and pyruvate dehydrogenase activity. Moreover, we determined that the metabolic derangements in the absence of FrmB impact formation of small colony variants (SCVs), a clinically relevant metabolic transition. Importantly, *S. aureus* does not demonstrate SCV-linked resistance to a known prodrug, POM-HEX, and loss of FrmB resensitizes *S. aureus* to this prodrug under SCV-inducing conditions.

Our data strongly suggest that FrmB controls steady state levels of pyruvate in staphylococci. Prior evidence indicates that pyruvate plays a central role in the modulation of *S. aureus* virulence; however, most of this work has been based on exogenous addition or environmental exposure to pyruvate. For example, upon pyruvate exposure, *S. aureus* induces the production of cellular toxins called leucocidins in an AgrAC-, SaeRS-, and ArlRS-dependent manner, thus enhancing virulence.^34^ Exogenous exposure to pyruvate has been linked to an increased production of acetate by *S. aureus*, which corresponds to increased cell viability under highly acidic environments, as can be found in abscess fluid or intracellular lysosomes.^35^

While prior evidence has established that exogenous pyruvate impacts the ability of *S. aureus* to infect and survive, our work demonstrates that perturbations in *intracellular* pyruvate levels and pyruvate dehydrogenase (PDHC) activity also impact the fitness of *S. aureus.* Pyruvate homeostasis has also been implicated in pathogenesis of *S. aureus* in an osteomyelitis mouse model.^36^ Strains lacking enzymes involved in pyruvate metabolism, such as pyruvate kinase (*pyk)* or the E1 subunit of PDHC *(pdhA)* are attenuated in survival during bone infection, suggesting that FrmB mediated regulation of PDHC may also be important in the context of bone tissue infection (supplemental figure 3). Interestingly, the attenuation phenotype this group established appears bi-modal in the context of *pdhA*, which they posit may be due to alternate fates of pyruvate, such as pyruvate being converted directly into oxaloacetate by *pyc* rather than *pdhA* mediated conversion into acetyl-CoA (supplemental figure 3).Their hypothesis may also explain our observations with respect to *S. aureus* TCA cycle intermediate levels when FrmB is not present. Lacking FrmB resulted in a defined metabolic blockage upstream of pyruvate, but a non-significant trend downward with a larger margin of error with respect to TCA-cycle intermediates. It is possible that *S. aureus* compensates for a lack of PDHC efficiency and shunts pyruvate into the TCA cycle to ameliorate pyruvate accumulation.

We observed that FrmB impacts the native function of PDHC, but does not impact the total protein levels of PDHC, suggesting that FrmB may interact with PDHC or PDHC related factors. Our study assessed PDHC function utilizing an assay that measures the E1 subunit of PDHC, suggesting that the level of regulation conferred by FrmB occurs at the E1 subunit, or with known E1 subunit factors. In the well-studied mammalian PDHC, the E1 subunit is subjected to phospho-regulation by a large number of dedicated kinases and phosphatases.^37^ However, prokaryotic regulation of the E1-PDCH subunit is poorly understood and no PDHC kinases or phosphatases have been robustly classified. Still, a study in *Corynebacterium* spp. identified potential acyl modification sites in multiple central metabolic enzymes, including the *Corynebacterium* E1-PDHC, suggesting possible post-translational modification sites.^38^ It is possible that the E1-PDHC in *S. aureus* is likewise regulated through post-translational modification, and future studies will evaluate the impact of FrmB on these PTMs.

While the *S. aureus* E1 sub-unit is relatively understudied, *S. aureus* PDHC-E2 regulation has been well characterized.^39–42^ PDHC-E2 is one of four enzymes in *S. aureus* that are lipoylated, required for proper functioning of the enzyme. Either by scavenging or *de novo* synthesis of lipoic acid, *S. aureus* decorates the E2 subunit of PDH with a lipoyl group via the amidotransferase LipL. Lack of lipoylation of the E2-subunit of PDHC renders the PDHC complex defective. The E2 subunit interacts with the E1 subunit by its lipoyl cofactor accepting the acyl group of the decarboxylated pyruvate and subsequently transferring the acyl group to coenzyme A. ^39–42^ In this study, we observe that FrmB impacts PDHC function by measuring PDHC E1-mediated decarboxylation. However, it is possible that FrmB does not directly impact the E1 subunit of PDHC, but instead interacts with the E2 subunit. If FrmB interacts with the lipoyl based modification pathway of PDHC E2, lacking FrmB might hinder the acyl transfer from E1 to E2 and impact the upstream E1 decarboxylation step our assay measures.

Small colony variants (SCVs) represent a significant challenge for clinicians due to their enhanced antibiotic resistance garnered by metabolic quiescence.^19–20^ Previous work has indicated that small colony variants decrease cellular respiration and upregulate glycolysis, pyruvate metabolism, and specific amino acid metabolism, and indeed, our results demonstrate that FrmB impacts the ability of *S. aureus* to form small colony variants, likely due to its involvement in central carbon metabolism.^20–24^ Our lab has previously identified FrmB as a staphylococcal esterase capable of activating a prodrug enolase inhibitor, POM-HEX.^25,44^ We know that resistance to this prodrug can be acquired via point-mutations in FrmB upon persistent POM-HEX exposure.^25,44^ However, our data from this study demonstrate that lacking FrmB *resensitizes S. aureus* to POM-HEX under SCV inducing conditions. This result has important implications with respect to structure-based prodrug design development focused around the FrmB esterase, suggesting that acquired mutations from POM-HEX exposure do indeed decrease the overall fitness of the bacteria, rendering FrmB a viable target for future prodrug’s designed with FrmB substrate specificity in mind.

Overall, our study bolsters the understanding of the role of intracellular pyruvate in *S. aureus* fitness and factors governing pyruvate metabolism. We demonstrate that a serine hydrolase, FrmB, contributes to the metabolic regulation of pyruvate and that lacking this enzyme result in fitness loss and the inability to transition into a small colony variant state. Importantly, we demonstrate that FrmB is a suitable target for future studies aimed at developing novel prodrugs based around FrmB substrate specificity. This work emphasizes the need to further understand *S. aureus* metabolism at all levels to combat *S. aureus* associated chronic disease states, infection, and persistence.

## Methods

### Strains used and growth conditions

A *Staphylococcus aureus* JE-2 background was utilized for this study. An FrmB transposon insertion mutant in the JE-2 background was leveraged (*frmB::*tn) to investigate the biological impacts of FrmB on *S. aureus*.^46^ For targeted metabolomics, bacterial cultures were grown in a complete M63 media (100mM potassium dihydrogen phosphate, 15.12mM ammonium sulfate, 0.0004mM ferric citrate, 16.7mM glucose, 0.409mM biotin, 0.0163mM nicotinic acid, 1mM magnesium sulfate, 1x EZ supplement from teknova, 1x ACGU supplement from teknova) with or without supplemented carbon sources and adjusted to a pH of 7.4. For nutrient add-back assays, cultures were grown in an RPMI SILAC flex media supplemented with L-arginine and L-lysine (supplementary table 1), but lacking major carbon sources required to support *S. aureus* growth. Nutrient add-back assays were conducted by adding 7.8mM of either glucose, sodium pyruvate, or sodium acetate. Bacterial growth curves were measured every 30 minutes at a wavelength of 600nm in a 96-well plate reader. Competition assays, small colony variant generation, and small colony variant persister assays were likewise completed in the above RPMI SILAC flex media supplemented with L-arginine and L-lysine, but included 7.8mM of glucose to support equivalent growth of all strains. Cultures, unless otherwise noted, were grown at 37°C, shaking at 250rpm.

All growth media ingredients and catalog numbers are additionally listed in supplementary table 1.

### Targeted metabolomics growth conditions

JE-2 and *frmB::*tn were both grown to an OD600 of 0.5 in M63 complete media in biological replicates of 5. All samples were subsequently pelleted and flash frozen. Samples were sent to Metabolon, Inc. on dry ice and processed using their Ultrahigh Performance Liquid Chromatography-Tandem Mass Spectroscopy (UPLC-MS/MS) platform. Sample processing parameters as performed by metabolite are described in Supplementary methods 1.

### Targeted metabolomics data analysis

#### Filtering and normalization of raw data

The Metabolon UPLC-MS/MS platform detected a total of 309 compounds from the panel: 259 compounds of known identity and 50 compounds of unknown identity. Initial filtering and normalization of raw data was performed in R using the metaboanalyst package: Raw peak values were first filtered (features with low repeatability as measure by relative standard deviations > 25%, and variables that are near constant throughout the experiment determined by IQR values, filtering out 10%.) Filtered raw peak values were then normalized based on DNA content for each sample and log transformed. Two samples (CHPH-03065 and CHPH-03088) were determined to be statistical outliers and were omitted from statistical analyses. Statistical outliers were determined by comparing each sample to all the other samples within its group for every metabolite. If the sample was an outlier for >40% of the metabolites, the sample was deemed an outlier. Metabolite outliers were determined by subtracting 1.5 x the interquartile range (IQR) from the first quantile and adding 1.5 x the IQR to the third quantile. Values outside this range would be considered outliers. Raw targeted metabolomics data is included as supplementary table 2.

#### Statistical analyses

All downstream analyses were also performed in R.

PCA plots were generated using the metaboanalyst package. A permutational multivariate analysis of variance (PERMANOVA) was calculated using Adonis2 function from the Vegan package. Distance matrices were calculated using the Eucledian dissimilarity metric on log-transformed and DNA-normalized data. Group differences were assessed using 10,000 permutations. A hierarchical clustering analysis was performed using the pheatmap package. The data represented are from normalized values with features auto-scaled, ward clustering, and Euclidean distance measured. The top 50 samples based on Z-score are presented. Fold change was calculated on normalized values and significant values were determined by t-test. A fold change threshold of 2.0 and a p-value threshold of 0.05 were applied. A volcano plot was visualized using the ggplot2 package. Box plots depict normalized abundance values for each feature, statistical significance determined by t-test. Box plots were generated using the boxplot function in R. Pathway enrichment was performed utilizing metaboanalyst package and visualized using ggplot2. Global ANCOVA enrichment analysis was performed on the normalized metabolomics data The topology measure used was relative-betweenness centrality and the Staphylococcus aureus subsp. aureus USA300FPR3757 (CA-MRSA) (KEGG) metabolome was used as the reference. Pathway impact is reported against statistical significance. False discovery rate (FDR) is also reported.

### FrmB complementation

Complementation of FrmB was performed as previously described.^45^ The complete coding sequence of *frmB* and 100 base-pairs upstream from the *S. aureus* JE-2 genome were cloned as a single copy into the *S. aureus* pathogenicity island 1 (SAPI1) by phage transduction.

Confirmation of *frmB* insertion was performed via PCR using *frmB* specific primers. Complementation of FrmB activity was performed utilizing an established fluorescent ester-based enzyme assay.^25^ FrmB has been previously shown to have high specificity for an ester 6C. Whole cell bacterial lysates of JE-2, *frmB*::tn, and the complement were added to a prewarmed buffer (25 mM Tris pH 7.5, 250 mM NaCl, 1 mM MgCl2, 10% glycerol) at a final concentration of 30 µg/mL of total protein. Enzyme catalyzed substrate hydrolysis was initiated by the addition of 5 µL of a 500uM stock ester 6C. The resulting change in fluorescence (λex = 485 nm, λem = 520 nm) was followed for 15 minutes at 37°C, collecting data every 30 seconds on a FLUOstar Omega microplate reader (BMG Labtech). Fluorescent ester activation was utilized as an indicator of FrmB functional restoration. Two way-ANOVA with Tukey’s correction for multiple comparisons was performed using GraphPad Prism version 10.0.0 for Windows, www.graphpad.com.

### Pyruvate Dehydrogenase (PDHC) Activity Assay

Bacterial whole cell lysates were prepared by growing each strain to mid-exponential phase in RPMI SILAC flex media supplemented with amino acids and glucose necessary to support equivalent growth (supplemental table 2.) Cultures were spun down at 10,000rpm and resuspended in Dulbecco’s Phosphate-Buffered Saline (DPBS) for sonication. All isolates were kept on ice and immediately flash frozen in aliquots and stored in -20°C until further use. To assess pyruvate dehydrogenase (PDHC) activity, an established 2,6-Dichlorophenolindophenol (DCPIP) reduction assay was utilized, whereby the E1 subunit of PDHC reduces DCPIP and can be monitored for change in absorbance.^30^ 6 μg of whole cell bacterial lysate was added to 0.2mM thiamine pyrophosphate, 2mM magnesium chloride, 50uM DCPIP, and 100mM potassium phosphate (adjusted to a ph =7.0.) After a 10 minute incubation period at 30°C, the reaction was initiated by the addition of 100mM sodium pyruvate. The rate of DCPIP reduction was calculated by monitoring the change in optical density over time at 30°C at a 600nm wavelength and using the DCPIP absorption constant of ε_600_ = 11,000 M^−1^ cm^−1^. Statistical analysis was performed in Prism. Assays were performed in biological triplicate in technical triplicate. One way-ANOVA with Tukey’s correction for multiple comparisons was performed using GraphPad Prism version 10.0.0 for Windows, www.graphpad.com.

### Western blots

Whole cell bacterial lysates utilized for the DCPIP activity assay were assessed for PDHC protein levels via western blot. 30 microliters of 6X sodium dodecyl sulfate (SDS) sample buffer (0.375 M Tris [pH 6.8], 12% SDS, 60% glycerol, 0.6 M dithiothreitol [DTT], 0.06% bromophenol blue) was added to the 90 μL of bacterial whole cell lysate, and the mixture was boiled for 10 min at 98°C. 10 μL of sample was loaded into a 12% poly-acrylamide gel (0.75mm) and allowed to run for ∼1 hour at 110 V. Transfer was performed using the Bio-rad instrumentation using home-made (cut) stacks and PVDF membrane.

Next, the membrane was transferred to 5% BSA in PBST and allowed to rock at room temperature for 1 hour. Rabbit polyclonal anti-lipoic acid antibody (Calbiochem) was applied as the primary antibody at a 1:7,500 dilution in cold blocking solution (5% BSA.) Rabbit polyclonal FtzS (MyBioSource, #<u>MBS540208</u>) was utilized as a loading control at a 1:3750 dilution in cold blocking solution (5% BSA.)

After 1 hour incubation at room temperature, the membrane was washed 3x for 10 minutes in cold PBST. Next, anti-Rabbit IgG from goat (HRP conjugated) was diluted 1:20,000 in fresh cold blocking solution (5% BSA) and added to the membrane. After a 1 hour incubation at room temperature, the membrane was washed 3x for 20 min in cold PBST and prepped for imaging.

### Bacterial competition

A 1:1 ratio of JE-2 and *frmB::*tn was added to 5mL culture tubes of RPMI SILAC flex media supplemented with amino acids and glucose necessary to support equivalent growth (supplemental table 2) and allowed to grow for 24 hours. At zero hours and 24 hours, culture was taken for serial dilutions and quantifying colony forming units (CFUs). Dilutions were plated on LB and LB plates treated with 5ug/ml of erythromycin (erm), from which a competitive index was calculated as follows: total CFUs on LB/erm5 divided by the total CFUs on plain LB. Because *frmB::*tn contains erythromycin resistance by virtue of the transposon insertion^44^, a competitive index of 0.5 would indicate no competitive difference when lacking a functional FrmB, an index >0.5 would indicate a competitive advantage when lacking a functional FrmB, and an index < 0.5 would indicate a competitive disadvantage when lacking a functional FrmB. Six biological replicates were performed in technical duplicate. Welch’s t-test was performed using GraphPad Prism version 10.0.0 for Windows, www.graphpad.com.

### Extracellular flux analysis

Extracellular flux analyses were conducted on the Agilent Seahorse Xfe96 Analyzer. Bacterial cultures were initially grown up to an OD600 of 0.5 in RPMI SILAC flex media supplemented with amino acids and glucose necessary to support equivalent growth (supplemental table 2.) Subsequently, each bacterial culture was spun down, washed in Agilent DMEM minimal media, and diluted back to an OD600 of 0.05 in Agilent DMEM for subsequent seeding on a poly-d-lysine coated seahorse compatible 96 well plate.

The 96 well plate and associated injection ports were placed in the instrumentation, and baseline ECAR and OCR levels were taken for each well for three cycles prior to carbon source injection. After baseline normalization, 10mM of each carbon source (glucose or pyruvate) was injected into each experimental well for a final carbon source concentration of 1mM per well.

Measurements were taken in 160 second cycles with gentle mixing prior to each reading. Each strain and condition was assayed in three technical replicates and three biological replicates. Two way-ANOVA with Tukey’s correction for multiple comparisons was performed using GraphPad Prism version 10.0.0 for Windows, www.graphpad.com.

### Small colony variant generation and persister assays

Small colony variants (SCVs) were generated as previously described.^31^ Briefly, bacterial isolates were grown in RPMI SILAC flex media supplemented with L-arginine, L-lysine, and glucose. The media was adjusted to a pH of 7.4 or 4.0 using citric acid. Isolates were allowed to grow for 24 hours statically at 37C. After 24 hours, CFUs and SCVs were enumerated by serial dilution on LB plates. SCVs were defined as pin-point colonies 1/10^th^ the size of a normal *S. aureus* CFU.

SCV persister assays and IC50s were set up as previously described.^31,33^ All isolates were grown in the same manner as the SCV generation protocol above, but at 24 hours, each culture was spun down, washed, and resuspended in fresh RPMI SILAC flex media supplemented with L-arginine, L-lysine, and glucose adjusted to a pH of 7.4. Immediately, antibiotic IC50s were set up using these washed cultures in the same media adjusted to a pH of 7.4. Each assay was set up as three biological replicates with technical duplicates. Two way-ANOVA with Tukey’s correction for multiple comparisons was performed using GraphPad Prism version 10.0.0 for Windows, www.graphpad.com.

### Bacterial inhibition (IC50s)

Bacterial IC50 assays were performed using microtiter broth dilution in clear 96-well plates. 10 serial dilutions were performed for each compound in technical duplicates for three biological replicates. A well containing bacteria but no drug was used as a positive control, and a well containing LB and no bacteria was utilized as a negative control for contamination. A starting concentration of 200uM (for conditions pre-grown in a pH of 7.4) or 40uM (for conditions pre-grown in a pH of 4.0) of POM-HEX was utilized. Plates were inoculated with 75 μL of bacteria diluted to 1 × 10^5 CFU/mL in LB. After inoculation, plates were incubated for 16 hours while shaking at 250 rpm and 37 °C. IC50 curves were calculated in prism by applying a non-linear regression dose-response inhibition curve (variable slope, four parameters) to log transformed and normalized values. Mean IC50 values are reported for each biological replicate, and a representative curve is provided for each condition. IC50s were conducted as three biological replicates with two technical replicates. Two way-ANOVA with Tukey’s correction for multiple comparisons was performed using GraphPad Prism version 10.0.0 for Windows, www.graphpad.com.

## Data availability

The data that supports the findings of this study are openly available in the supplemental materials (metabolomics raw data provided in supplemental table 2) or upon request from the corresponding author.

## Acknowledgements

This work was made possible by funding from the National Institute of Health (NIH/NICHD R01HD109963)(NIH/NIAID R01AI171514) and the Microbial Pathogenesis and Genomics T32 Program at the University of Pennsylvania (5 T32 AI 141393-05). We are grateful to Florian Muller for the synthesis and provision of Pom-HEX, Paul Planet for providing reagents and necessary vectors for complementation construction, and to the lab of Douglas C. Wallace for the training and utilization of their Agilent Seahorse instrumentation.

